# A state-dependent mean-field formalism to model different activity states in conductance based networks of spiking neurons

**DOI:** 10.1101/565127

**Authors:** Cristiano Capone, Matteo di Volo, Alberto Romagnoni, Maurizio Mattia, Alain Destexhe

## Abstract

Higher and higher interest has been shown in the recent years to large scale spiking simulations of cerebral neuronal networks, coming both from the presence of high performance computers and increasing details in the experimental observations. In this context it is important to understand how population dynamics are generated by the designed parameters of the networks, that is the question addressed by mean field theories. Despite analytic solutions for the mean field dynamics has already been proposed generally for current based neurons (CUBA), the same for more realistic neural properties, such as conductance based (COBA) network of adaptive exponential neurons (AdEx), a complete analytic model has not been achieved yet. Here, we propose a novel principled approach to map a COBA on a CUBA. Such approach provides a state-dependent approximation capable to reliably predict the firing rate properties of an AdEx neuron with non-instantaneous COBA integration. We also applied our theory to population dynamics, predicting the dynamical properties of the network in very different regimes, such as asynchronous irregular (AI) and synchronous irregular (SI) (slow oscillations, SO).

This results show that a state-dependent approximation can be successfully introduced in order to take into account the subtle effects of COBA integration and to deal with a theory capable to correctly predicts the activity in regimes of alternating states like slow oscillations.

## Introduction

Recent developments in recording techniques are shading light on the dynamics of cortical neural networks in higher and higher spatio-temporal detail [1]. There are different scientific ways to investigate and understand such large amount of data. A first class of approaches are top-down aiming to use data as a constrain to build generative models capable to automatically reproduce statistical features observed in experiments. [2, 3, 4, 5]. On the other hand it is possible to interpret the experimental observed behavior by mean of a bottom-up theoretical model. To achieve this different levels of description are possible, ranging from single spiking neurons [6, 7] to population model [8, 9, 10, 11, 12, 13, 14, 15], from extremely detailed [16, 17, 8, 18] to more coarse-grained models [19, 20].

While keeping the model as simple as possible, it has been recently showed that some minimal requirements are necessary in order to reproduce a rich repertoire of observed features. In particular, a quite refined model as the AdEx is necessary to describe a response on a broad range of frequencies [21]. Moreover, voltage dependent synapses have been largely shown to be a crucial mechanism of neuron interactions [22]. While direct simulation of large ensembles of units can be performed, such an approach can be computationally heavy and do not permit a straightforward understanding of the system dynamics. A principled dimensional reduction approach such as mean-field theories (MF) are powerful and widespread tools, used to obtain large scale description of neuronal populations. One of the first successful attempt was to provide a theory to describe leaky integrate and fire neurons with current based input [19, 23], where the firing rate properties of neurons are described as a function of the infinitesimal average and variance of its input current by the use of a Fokker-Planck formalism. Also the effect of synaptic integration could be considered [24] that has the net effect to move the neuronal firing threshold. Later on, various attempts have been performed in order to extend this theory to voltage dependent synapses. After approximated approaches [25], an exact solutions has been formalized [26]. While such theory provides a good estimation in the case of delta-like synapses, the formalism does not provide an explicit solution in the case of synaptic integration.

Indeed to take into account these features all together is very challenging and despite a semi-analytic approach can give satisfactory quantitative predictions [27, 28] a simple and closed analytic solution has not yet been addressed.

Here we show how different approximations can lead to two different analytic results and how each of the two approximation only work in a specific dynamical condition, that can be either drift driven or fluctuation driven. For this reason we proposed a principled state-dependent approximation. In other words we showed that the two approximations are valid in the two limit described above and that they can be analytically merged. This allows to define a current-to-rate gain function reliable also in regimes where the dynamics is not strictly drift or fluctuation driven.

Our approach turns out to be particularly powerful when applied to investigate the properties of neural populations dynamics. In particular we considered a network composed of excitatory and inhibitory neurons, that is considered to be the standard minimal circuitry for cortical neuronal networks [29, 30]. The network parameters are set reproduce two different dynamical condition that are biologically relevant, i.e. asynchronous irregular and slow oscillating dynamics. We show that both of them are reliably described by our mean field model and that the state-dependent approach is indispensable to achieve the quality of such result.

## Results

### Neuronal network model

We derive a state-dependent current-to-rate gain function for conductance based (COBA) AdEx type neurons, whose dynamics evolve according to the following equations [7]:

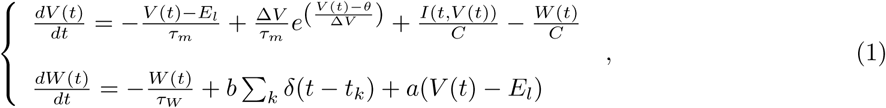

where the synaptic input *I* is defined as

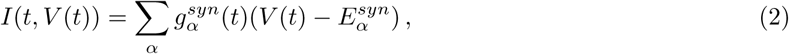

*V* (*t*) is the membrane potential of the neuron and *α* = *e, i* defines the excitatory (*e*) and the inhibitory (*i*) input. The population dependent parameters are: *τ*_*m*_ the membrane time constant, *C* the membrane capacitance, *E*_*l*_ the reversal potential, *θ* the threshold, Δ*V* the exponential slope parameter, *W* the adaptation variable, *a* and *b* are the adaptation parameters, and 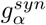 the synaptic conductance, defined as

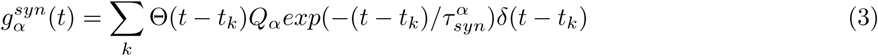

We define the spiking time of the neuron when the membrane potential reaches the threshold *V*_spike_ = *θ* + 5Δ*V*. 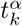 indicates the times of pre-synaptic spikes received by the neuron from synapse type *α* with characteristic time 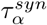 and its synaptic efficacy *Q*_*α*_.

### Current-to-rate gain function

Under the assumption of quasi-instantaneous synaptic transmission (negligible 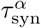), for a neuron described by the dynamical system of Eq. (1) it is possible to write a Fokker-Planck equation describing the dynamics of the probability density function (p.d.f.) for its membrane potential *V* as

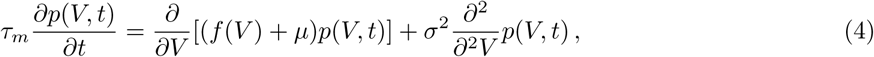

where we assumed that the input 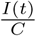 is a white noise with mean *μ* and variance *σ* (Fig1.A), 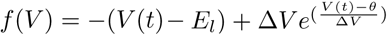 and suited boundary conditions are taken into account [31], i.e. an absorbing barrier at the spike emission threshold *V*_*spike*_ and that the probability current 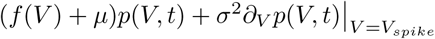 is re-injected in *V*_*reset*_ = − 65 after a refractory period *τ*_*arp*_ = 5*ms*. Under stationary conditions, the firing rate of the neuron is given by the flux of realizations (i.e., the probability current) crossing the threshold *V*_*spike*_ [23]:

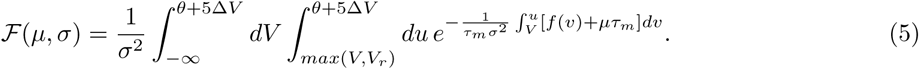

Such function, usually referred to as transfer function (or current-to-rate gain function), provides an estimate of neuronal firing rate which is in remarkable agreement with the one measured from numerical integration of Eq. (1) (Fig. 1B). Nevertheless, in the case of voltage dependent synapses determining the infinitesimal moments of the input current (mean *μ* and variance *σ*^2^) as a function of the input firing rate is not straightforward.

**Figure 1:**
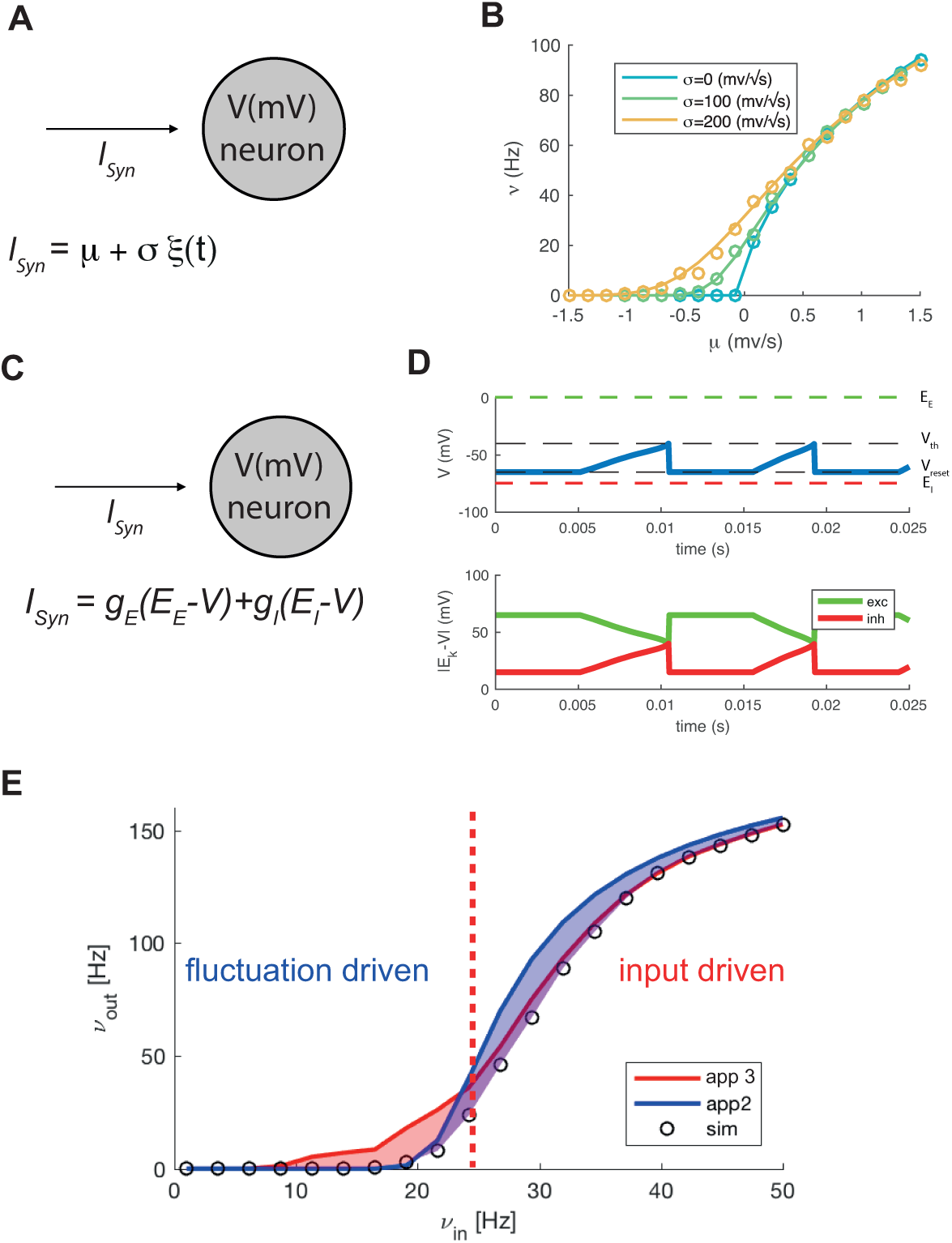
Current-to-rate gain function for AdEx neurons with conductance based input: (A) Sketch of a AdEx neuron with current based input represented by a white noise. (B) Current-to-rate gain function ℱ(*μ, σ*) for AdEx neuron receiving a white noise input with mean and variance (*μ* and *σ*, respectively). Theory and simulations (lines and circles, respectively) are in remarkable agreement. (C) Sketch of an AdEx neuron with conductance-based (COBA) input. (D) Graphic presentation of the voltage dependence of the conductance based input. (E) Firing rate of neuron with COBA input as a function of the excitatory input and with constant inhibitory one (circles). Two different theoretical approximations (in red and blue).

In particular when a conductance-based input is considered (Fig. 1C), the stochastic process describing the input current has voltage-dependent infinitesimal mean and variance due to the voltage-dependent nature of the impact of the incoming spikes on the membrane potential dynamics (Fig. 1D). In this framework, an explicit solution of the aforementioned Fokker-Planck equation has not yet been worked out.

### Moment Closure (MC) approximation

One of the major problems in modeling COBA neurons is that the input current is voltage dependent and can be written as

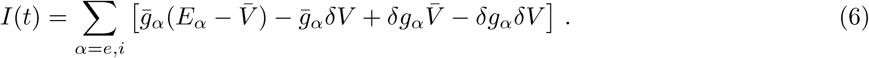

A first naive approximation consists in approximating the variable *V* by its average value 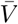, such that the input current 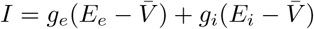 is now independent from *V*. Under diffusion approximation (i.e. in the limit of small *g*_*i*_ and *g*_*e*_, and a large rate of incoming spikes), the two infinitesimal moments *μ* and *σ*^2^ of *I* are:

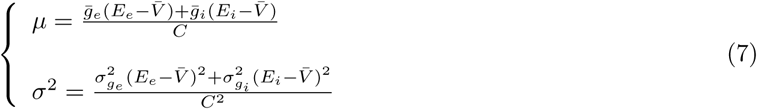

Since *μ* and *σ* can be written as a function of the firing rate, it is possible to write the transfer function

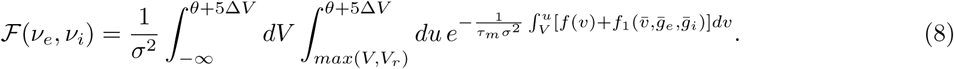

This equation is the same as Eq. (5) where 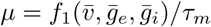 due to Eq. (7). Comparing this expression with numerical simulations of the single-neuron spiking activity in Fig. 1E (red line), a good agreement is mainly apparent under drift-driven regime (*μ τ*_*m*_ > *V*_spike_).

### Voltage-Dependent (VD) approximation

It is also possible to take into account the dependence of the input current *I*(*t*) on the voltage [26, 32, 33] by writing it in the following way:

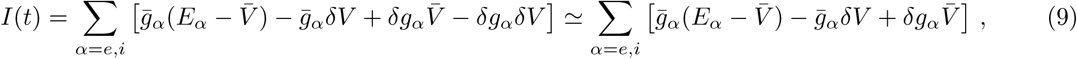

where we decomposed 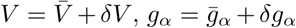 and the bar means the average over time. In the last step the term 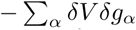 has been neglected since *δV* is assumed to be of the same order as 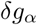 [26, 32], *so* 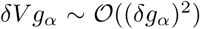. Under this approximation, the synaptic current can be then written as a deterministic voltage-dependent part plus a stochastic component which is independent from *V*. As we are considering a quasi-instantaneous synaptic transmission (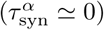), such stochastic source of current can still be modeled by a Gaussian white noise [23] such that:

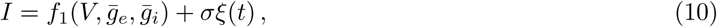

where 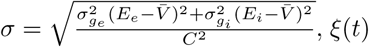 is a white noise *N*(0, 1) and 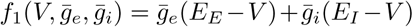, with 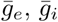 and 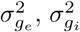 the mean and the variance of the synaptic conductances, respectively. In the case of input spike trains with Poissonian statistics these infinitesimal moments result to be [34, 23, 35]:

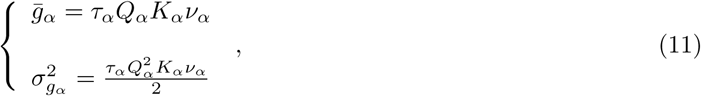

where *K*_*α*_ is the number of synaptic contact each neuron receives from the population *α* ∈ {*e, i*}.

As above, considering *f*_1_(*V*) as an additional term to *f* (*V*), it is again possible to work out an analytical expression for the transfer function:

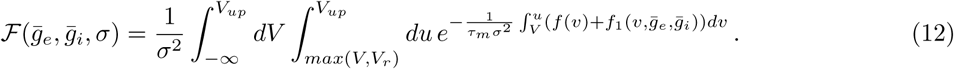

The result of such approximation is shown in Fig. 1E (blue line). We observe that this approximation gives good theoretical prediction as far as the average membrane potential of the neuron is sufficiently low (i.e. under noise-dominated regime).

### A mixed framework: State-dependent (SD) approximation

The previous two proposed approximations rely on different assumptions on the composition of the input current *I*(*t*) to the neurons, that turned out to be valid under different working regimes of the neuron. In this paragraph we propose a mixed framework in order to have a continuous transfer function, by introducing a new parameter that allows to interpolate between the two regimes. This parameter is introduced not by an *a posteriori* fit, but by *a priori* considerations on the input current.

Under drift-dominated regime (*μ τ*_*m*_ > *θ*), the spiking times are mainly determined by the deterministic component of the input and not by the stochastic one.

Accordingly, neglecting *V* fluctuations and replacing it with its average value, is a good assumption and the use of MC approximation is very satisfactory (Fig.1E left side).

When *μ τ*_*m*_ < *θ*, i.e. under *fluctuation-driven* regime, the neuron can only fire in presence of large-enough sub-threshold fluctuations, as 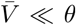. Therefore, all the variability of *V* has to be taken into account, as sub-threshold suppression appears when *V* is close to the *θ*. Under this condition, VD approximation result to be the most effective (Fig. 1E-right), as the additional term 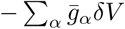 in the current *I*(*t*) is also taken into account. This term is lacking in the MC approximation.

Starting from that, we unify these two expressions for ℱ by writing 

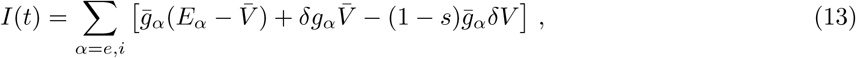

where *s* is an arbitrary state-dependent parameter which is 0 when 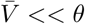 and 1 when 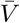 approached *θ*

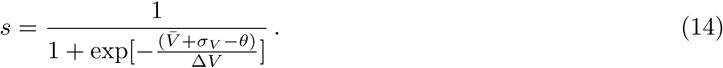

Here, *σ*_*V*_ and Δ_*V*_ are used because representing the natural scales for the system giving rise to following current-to-rate gain function:

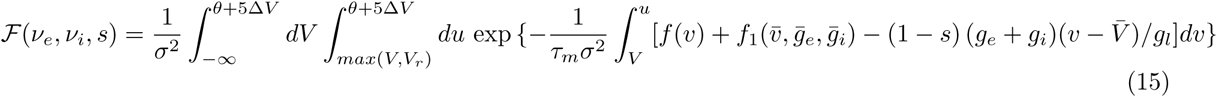

This formulation is valid in absence of synaptic integration 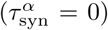, but its firing rate estimation is rather accurate even in presence of coloured input, as expected according to [24], as show in Fig. 2.

**Figure 2:**
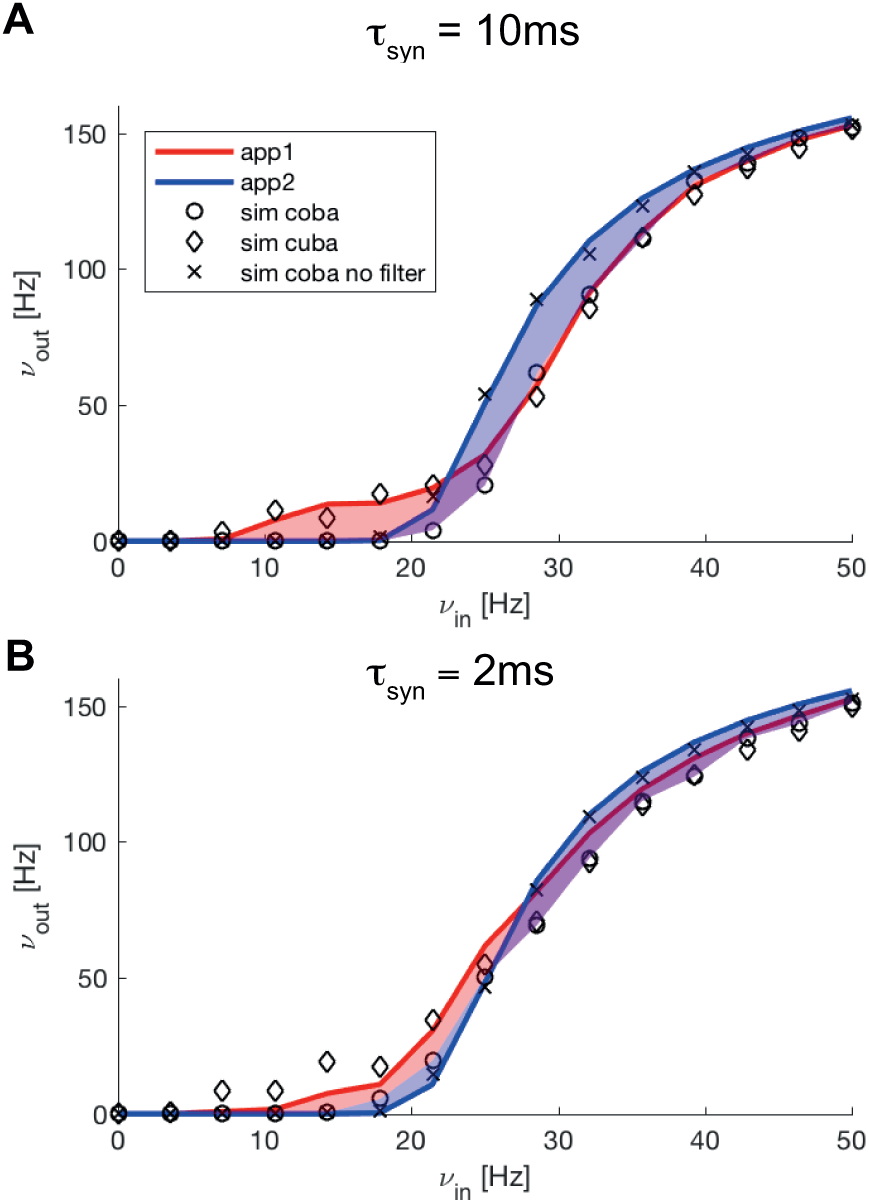
Different scales of synaptic integration. Comparison between two different scales of synaptic integration 5ms (A) and 1ms(B). Circles and crosses are COBA simulations respectively with and without synaptic filter. The diamonds are CUBA simulations. Approximation 1 (red) and 2 (blue) fit almost exactly CUBA simulations with synaptic filter and COBA simulations without synaptic filter.

In order to check the effectiveness of Eq. (15), we compared the ℱ obtained with the MC and the VD approximation, and with the state-dependent one, for varying excitatory and inhibitory input firing rates (Fig. 3A). We report the respective errors (difference between theory and simulations, see Fig. 3B) showing that in our approach they are smaller and distributed in a narrower region in the *v*_*e*_, *v*_*i*_ plane.

**Figure 3:**
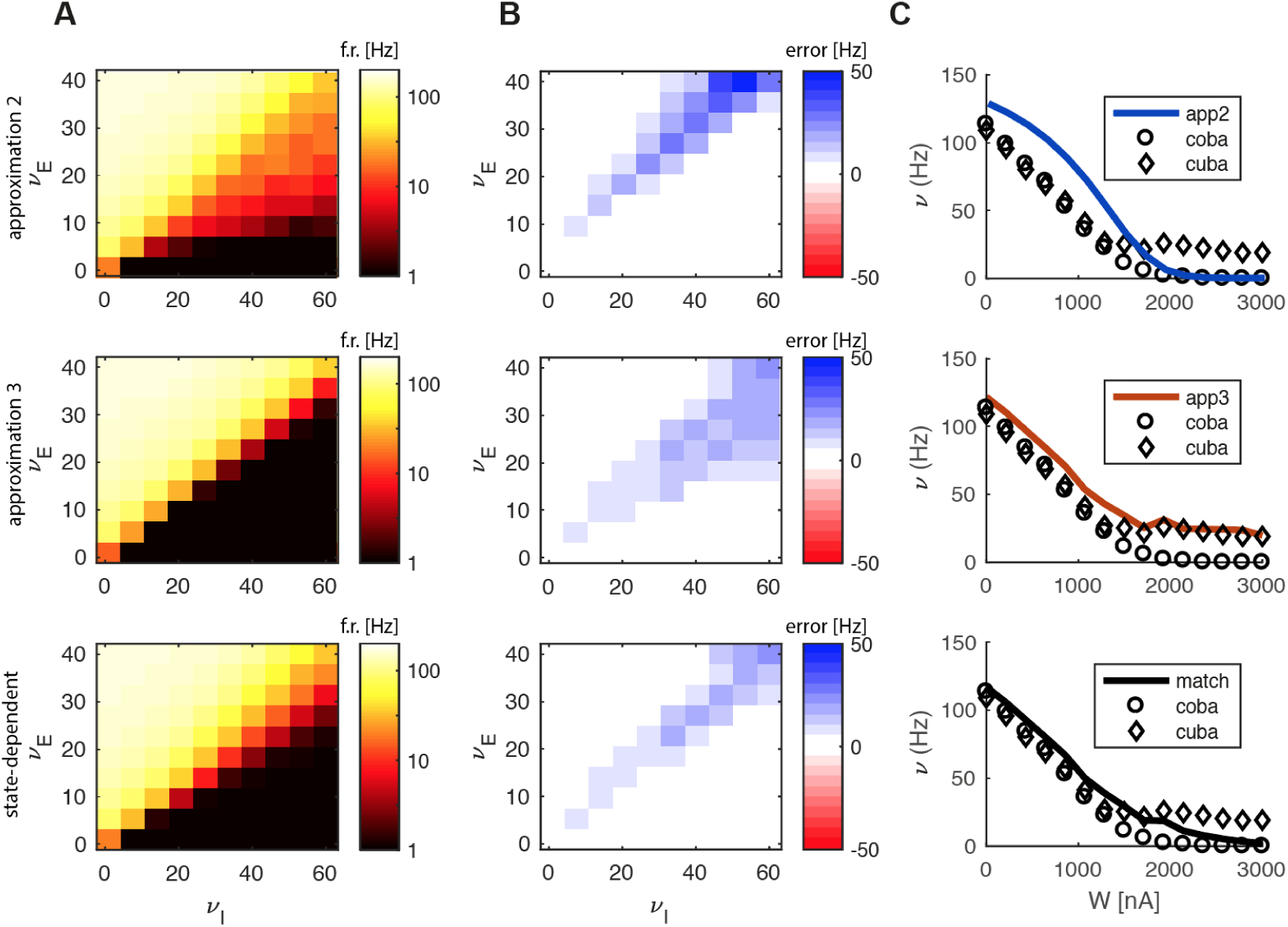
State dependent mean field approximation. (A) Theoretical predicted firing rate (color-coded) for the approximation 1, approximation 2 and the matched model. (B) Difference between the three theoretical models and the firing rate estimated in simulations. (C) Theoretical predicted firing rate (solid line) and firing rate from simulations for COBA and CUBA (respectively circles and diamonds) for the 3 theoretical models.

We considered also the adaptation variable *W* (*t*) with a relaxation time scale *τ*_*W*_ = *XXX* ms, and compared the prediction with the simulations with the three models, observing an optimal estimation for the state-dependent one (Fig. 3C).

### Application: population dynamics

We applied our result to describe an effective mean field dynamics for the canonically considered minimal structure of a cortical network, namely two coupled population of excitatory (regular spiking, RS) and inhibitory (fast spiking, FS) neurons, with spike frequency adaptation for the first type (see Fig.4A). The external input is provided by increasing the excitatory firing rate in the input of both the population by an amount of *v*_*ext*_ = 6*Hz*. Neuronal parameters are specified in Table 1. The probability of connection is *p* = 0.25.

**Table 1:** Neuronal parameters defining the two populations RS-FS model.

| | $\theta$<br>(mV) | $\tau_m$<br>(ms) | $C$<br>(nF) | $E_l$<br>(mV) | $\Delta V$<br>(mV) | $\tau_i^{syn}$<br>(ms) | $E_i^{syn}$<br>(mV) | $Q_i^{syn}$<br>(nS) | $b$ (nA) | $\tau_W$ (s) |
| --- | --- | --- | --- | --- | --- | --- | --- | --- | --- | --- |
| RS | -50 | 20 | 0.2 | -65 | 2.0 | 5 | 0 | 1 | 0.005 | 0.5 |
| FS | -50 | 20 | 0.2 | -65 | 0.5 | 5 | -80 | 5 | 0 | 0.5 |

We define the MF dynamics following the approach used in [36]

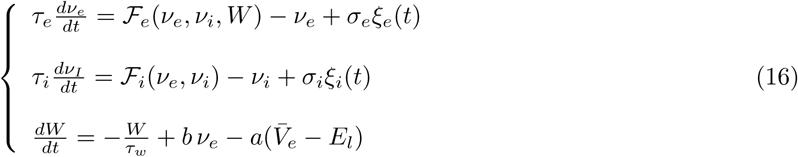

where we also considered the adaptation variable *W*. The parameter *b* and *a* are the same as in eq.(1). *τ*_*e*_ and *τ*_*i*_ are the same as the membrane potential time scales. *ξ*_*α*_ are white normal noises, and *σ*_*α*_ are the extents of the noise. 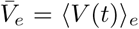 is the population average membrane potential. This is evaluated by integrating its deterministic differential equation. The adaptation corresponds to an additional term in the first infinitesimal moment, so that we can define

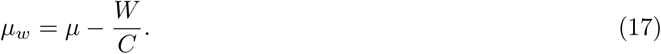

By changing the parameters, it is possible to set the network in different dynamical states. The asynchronous irregular (AI) is obtained by the parameters defined above. The slow oscillations (SO) are achieved by multiplying the probability of connection between excitatory neurons by a factor 1.15, increasing the excitatory adaptation strength to *b* = 0.02*nA* and decreasing the external input to *v*_*ext*_ = 0.95*Hz*.

The different regimes can be studied by the means of standard techniques used in dynamical systems theory, e.g. null-clines representation (see Fig.4B). Each null-cline represent the region where the derivative is zero for a certain variable (respectively blue for *v*_*e*_ and orange for *W*), and the intersection between them is a fixed point that can be either stable or unstable. The green line represents the dynamics in the plane (*v*_*e*_, *W*). This analysis is performed for the different choices of parameters and thus for the different dynamical conditions AI and SO. In panel Fig.4C is reported an example of the time-course of the dynamics for the two regimes (green and red respectively for *v*_*e*_ and *v*_*i*_). We eventually reported the average firing rate time-course for a network of spiking neurons with the same choice of parameters as in the previous analysis (Fig.4D), confirming that the predicted dynamics turns out to match the spiking simulations.

### Robustness of the prediction: need of a state-dependent approach

We tested the robustness of the prediction by exploring the parameter space. In particular we modified the external input in the AI, reporting the change in the stationary firing rates (Fig.5B), and the adaptation strength in the SO, reporting the Up and Down states duration (Fig.5D-E). The first two approximations taken alone poorly predicted the dynamics observed in the spiking simulations, while such task was performed quite well in the state-dependent approach.

**Figure 4:**
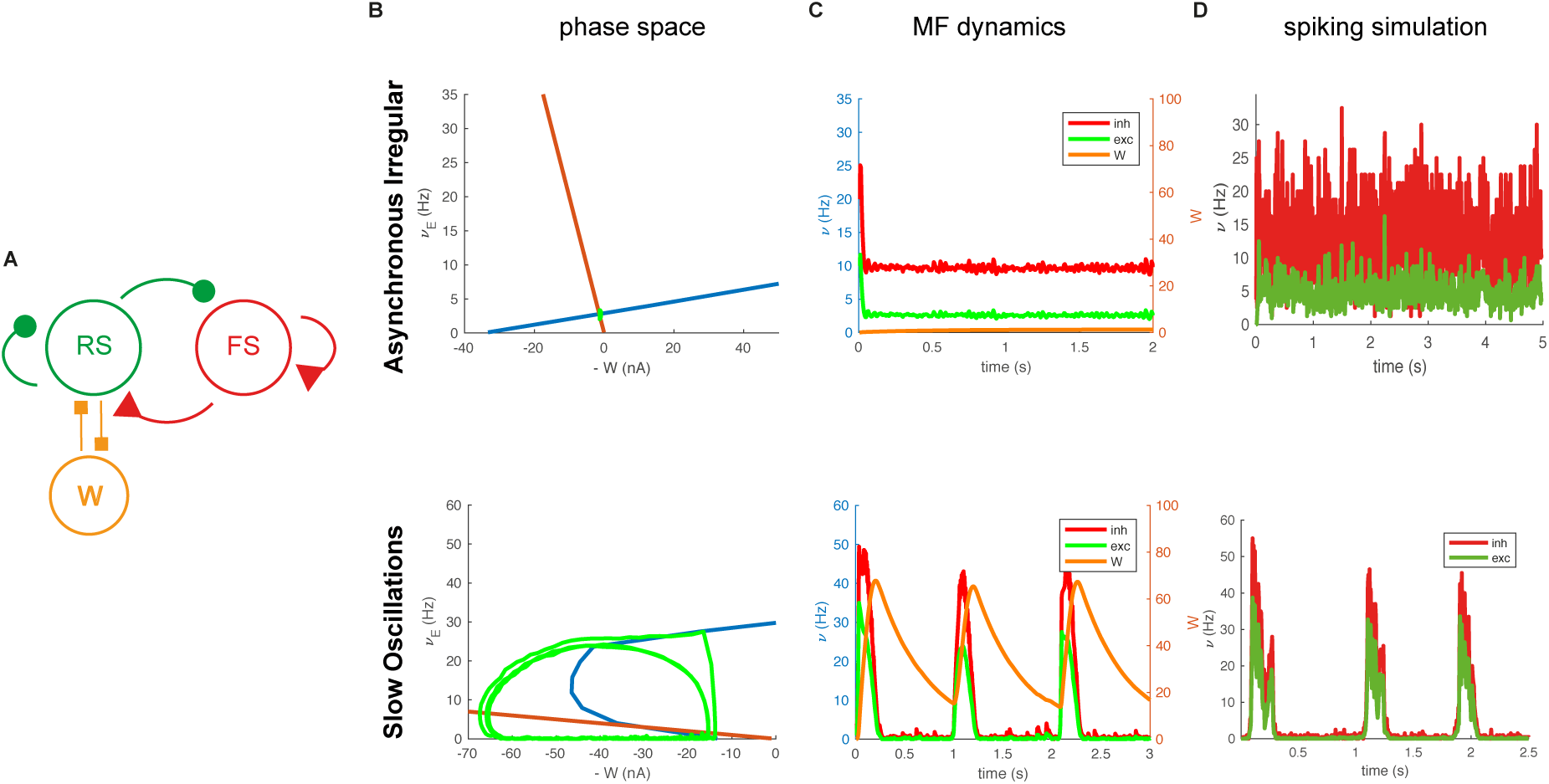
Mean field dynamics in a RS-FS network: (A) Sketch of the network structure. (B) Nullclines representation of the dynamical system in the phase space for 2 different dynamical regimes (Asynchronous Irregular and Slow oscillations). (blue and orange solid lines) Nullclines for the excitatory firing rate and the adaptation differential equation. (C) Example of mean field dynamics for the 2 different regimes (green and red represent excitatory and inhibitory firing rates respectively). (D) Average firing rate dynamics of the spiking simulation.

## Discussion

The mean field description of a large network of excitatory and inhibitory spiking neurons has been tackled analytically on relatively simple models, but often far from biophysical reality [19, 20]. On the other hand, anatomically sophisticated models [16, 17, 8, 18] are computationally consuming and very hard to be explored by mean of theoretical frameworks.

We proposed a tradeoff between these two possibilities. First we chose as neuron model, one of intermediate mathematical complexity yet high physiological validity, the exponential integrate-and-fire neuron with spike frequency adaptation. Second, we consider voltage dependent synapses (COBA) that so far made this problem difficult to be exactly solved.

To overcome the mathematical difficulty to solve a Fokker-Planck equation with a voltage dependent noise, describing a conductance based input, we proposed a mapping on a CUBA model, which has a known solution [19]. However, we showed that this mapping has to be state-dependent, since different approximations have to be considered in different regimes. However, we showed that this mapping relies on approximations that are state-dependent. Indeed, in the fluctuation-driven regime it is possible to use a standard approximation that basically maps the COBA on a CUBA with rescaled membrane time scale [26].

Nevertheless, in the drift-driven regime this approximation is no longer providing a good description, and it has been shown only to work in a relatively simple model with instantaneous synapses and leaky integrate and fire neuron. Our analysis reported that this is no longer valid when a synaptic integration is considered since this that creates a strong interaction between conductances and membrane potential. Nevertheless a different suitable approximation can be performed neglecting the fluctuations of the membrane potential, obtaining again an effective CUBA model where the variable the membrane potential *V* is frozen and replaced by a stochastic process with the same statistical moments. An analytic merge of the two approaches provides a good prediction of the firing rate in the whole phase space.

**Figure 5:**
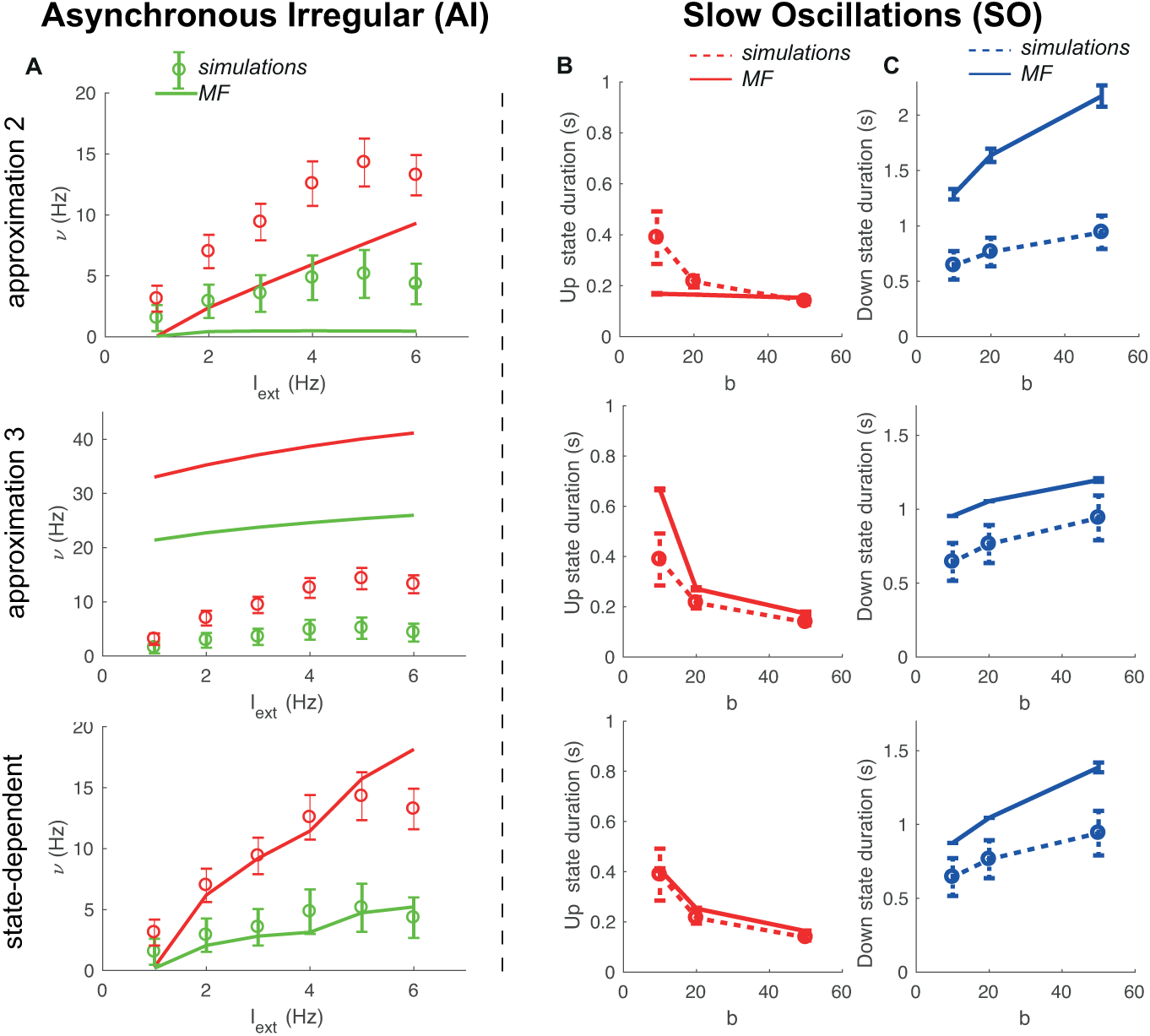
State dependent approximation is required to correctly capture the dynamics. (A) Predicted stationary excitatory and inhibitory firing rates (green and red lines) as a function of the amount of external noise, compared with spiking simulations (circles). (B-C) Predicted Up and Down states durations (solid line) as a function of the adaptation strength *b* compared with spiking simulations (dashed line).

Making approximations is a natural way to simplify a problem and understand more easily the underlying mechanisms. Our approach, since it relies on two different approximations, points out that the relevant aspects producing the observed dynamics are state-dependent.

Since neurons in cortical populations notoriously go across both noise and drift driven regime [37, 38, 39], to define a population mean field dynamics requires to take into account a unified framework like the one we proposed. To support this statement we have shown that when a single approximations have been considered the quality of predictions was extremely poor. A unique transfer function reliable in various dynamical conditions is particularly relevant also because different population may be in different regimes or the same population can change regime across time, as in the case of slow oscillations.

We showed that our method is robust and flexible and successfully describes different population dynamical regimes, such as asynchronous irregular state and slow oscillations.

Our approach suggests a general method to perform a state-dependent mapping of neurons with COBA input on to CUBA input even with different types of neuron such as QIF and LIF.

Our model could be interpreted as an attempt to do a step forward to the development of analytic but still rich and realistic theories that allow to describe experimentally observed phenomenons [22].

We propose that the model can be naturally extended to more complicated structures, such as the thalamo-cortical loop and network with spatial extension. This would permit to test our model on experimental data recording the activity of populations of neurons over space where it may provide a mechanistic understanding of the emerging dynamics based on neurons voltage based interactions.

A semi-analytic approach was proposed recently [27, 28]which relies on a fitting of the transfer function to numerical simulations. This approach yields mean field models of COBA neurons with good quantitative predictions. The main advantage provided by this ‘orthogonal’ approach is to be potentially applicable to any neuronal model and to experimental data. On the other side, as being a semiâanalytic fit, it does not permit the same understanding of the dynamical mechanisms underlying the neurons response function as a principled approach like it does the one here proposed. More detailed comparison of the two approaches is the object of future directions and the knowledge derived from these two different approaches will help to make important steps forward an unified theory about mean field models of COBA neurons.

## Acknowledgements

This project has received funding from the European Union’s Horizon 2020 Framework Programme for Research and Innovation under the Specific Grant Agreements No. 785907 (Human Brain Project SGA2) and No. 720270 (HBP SGA1).

